# Aging disrupts tissue homeostasis and constrains blastema-mediated regeneration in the *Cladonema* medusa

**DOI:** 10.64898/2026.01.21.700690

**Authors:** Ren Kanehisa, Hiroko Nakatani, Sho Takatori, Taisuke Tomita, Masayuki Miura, Yu-ichiro Nakajima

**Affiliations:** Laboratory of Genetics, Graduate School of Pharmaceutical Sciences, The University of Tokyo, Japan; Laboratory of Neuropathology and Neuroscience, Graduate School of Pharmaceutical Sciences, The University of Tokyo, Japan; Laboratory for Cell Vigor Regulation, National Institute for Basic Biology, Japan; Basic Biology Program, The Graduate University for Advanced Studies, SOKENDAI, Japan

## Abstract

Regenerative capacity varies widely across animals, yet aging is often accompanied by declining tissue homeostasis and regenerative potential. In many species, regeneration of complex structures relies on epimorphic programs that form a blastema, coordinating cell proliferation and pattern reorganization after injury. Although aging-associated regeneration defects have been documented in several bilaterian models, how aging shapes regeneration in early-branching metazoans with robust regenerative abilities remains unclear. Here, we investigate aging and regeneration in the medusa stage of *Cladonema pacificum*, a cnidarian that retains high regenerative capacity within a finite lifespan. We show that aging in *Cladonema* medusae is accompanied by progressive morphological and functional deterioration, including shrinkage of the umbrella and manubrium, tentacle shortening, and reduced reproductive output, along with increased DNA damage. At the cellular level, aging is associated with depletion of differentiated cell populations, including nematocytes and neurons, as well as reductions in stem cell-associated populations and proliferative activity in the tentacle bulb. Consistent with these changes, tentacle regeneration is markedly impaired in aged medusae and is characterized by delayed wound closure, defective blastema formation, and incomplete functional recovery. Together, our findings indicate that aging disrupts both tissue homeostasis and blastema-mediated regeneration in *Cladonema* medusae, establishing a tractable model for studying aging–regeneration interactions and supporting the view that aging imposes broadly shared constraints on regenerative systems across metazoans.

## Introduction

Aging is associated with a progressive decline in tissue homeostasis and regenerative capacity across metazoans. At the cellular level, aging influences multiple hallmarks, including stem cell exhaustion, altered intercellular communication, and loss of proteostasis, that together compromise tissue maintenance and repair (López-Otín et al. 2023). Consistent with this view, aging impairs stem cell function and regenerative responses across diverse tissues, leading to defective replacement of differentiated cells and reduced resilience to injury (Rando & Chang 2012; Goodell & Rando 2015). However, the cellular processes through which aging disrupts regeneration, particularly in animals with intrinsically strong regenerative capacity, remain incompletely defined.

Regeneration-competent models provide a powerful framework to dissect how aging interferes with tissue repair (Stoick-Cooper et al. 2007). In such systems, regenerative decline reflects aging-dependent alterations in cellular programs that normally support regeneration. Studies in regenerative vertebrates, for example, in the short-lived African turquoise killifish, aging impairs regeneration of complex tissues such as brain and fins by disrupting progenitor activation, proliferation, and early regenerative events (Wendler et al. 2015; Van houcke et al. 2021). Similarly, studies in axolotls have suggested that although regenerative capacity is retained throughout life, limb regeneration becomes progressively delayed with age, rather than being completely lost (Vieira et al. 2020; Faisal et al. 2024). These findings support the idea that aging acts as a conserved constraint on regeneration by targeting essential cellular contributors, yet whether similar principles apply to early-branching metazoans remains largely unexplored.

Cnidarians, as a sister group to bilaterians, occupy a key phylogenetic position for testing how age-associated changes influence regeneration (Fig. 1A). Despite their simple body organization, cnidarians possess conserved stem cell systems that support continuous tissue renewal and remarkable regenerative abilities (Bosch 2009; Holstein et al. 2003; Watanabe et al. 2009). Much of our understanding of aging in cnidarians derives from studies in *Hydra*, a sessile polyp that primarily reproduces asexually and can exhibit negligible senescence under laboratory conditions while maintaining stem cell function over extended periods (Martínez 1998; Schaible et al. 2015). In contrast, medusa-stage cnidarians are free-swimming and predominantly engage in sexual reproduction, representing a life-history stage with finite and physiologically constrained lifespans (Piraino et al. 2004). Thus, medusa-stage cnidarians provide a complementary system to examine how aging reshapes tissue homeostasis and regeneration. Nevertheless, how aging impacts tissue maintenance and regenerative capacity in cnidarian medusae remains poorly understood.

**Figure 1.**
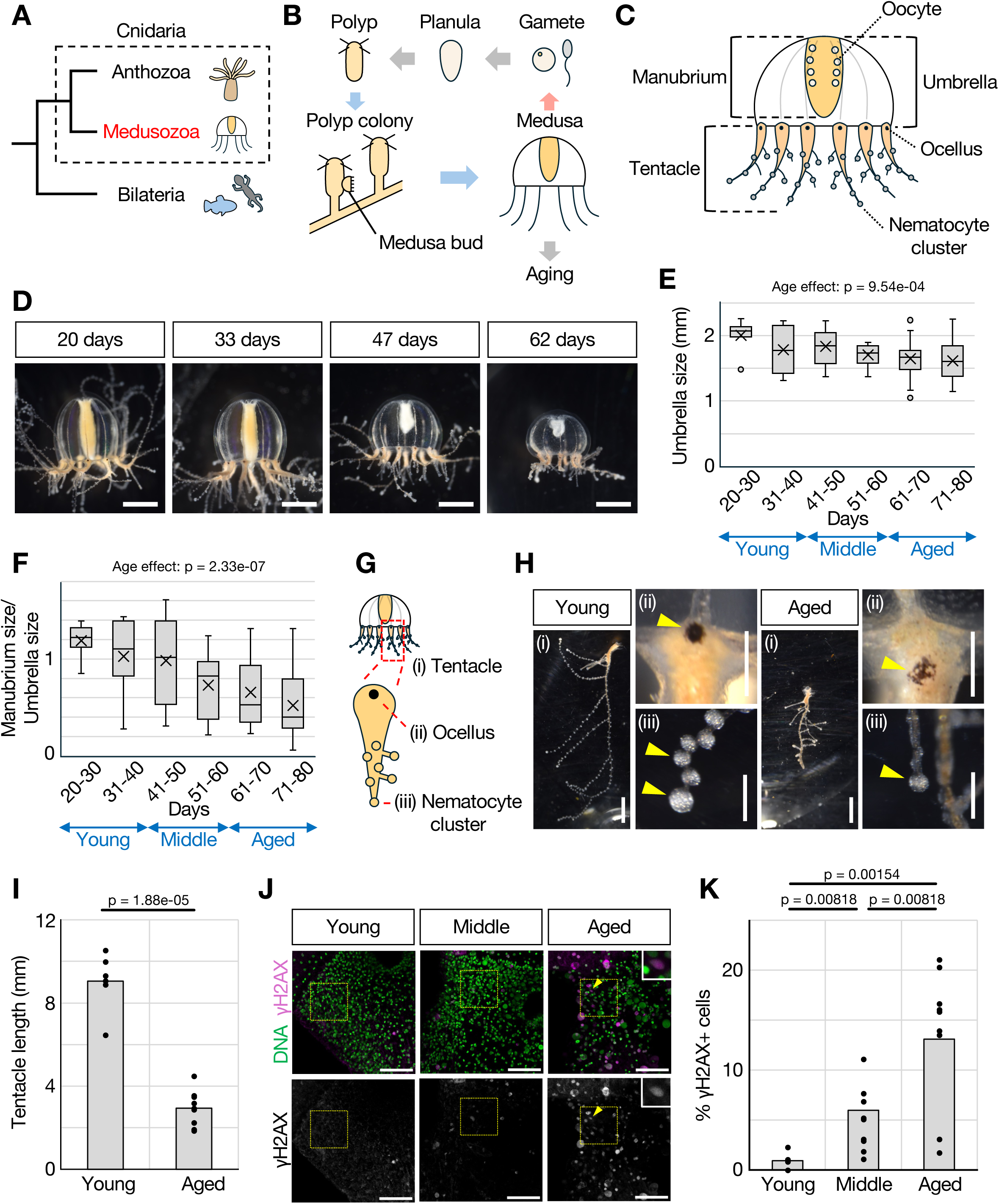
*Cladonema* medusae exhibit morphological and cellular changes with aging. (A) Phylogenetic relationship between Bilateria and Cnidaria. (B) Life cycle of *Cladonema pacificum*. (C) Schematic illustration of a female *Cladonema* medusa. (D) Representative images showing progressive age-associated morphological changes in *Cladonema* medusa. (E) Quantification of umbrella size showing an age-related reduction. Days 20-30: N=11 medusae, 31-40: N=7, 41-50: N=25, 51-60: N=11, 61-70: N=34, 71-80: N=53. Some individual medusae were longitudinally measured at multiple ages. The *p*-value indicates the overall effect of age assessed using a linear mixed-effects model. (F) Quantification of the ratio of manubrium size to umbrella size, showing relative shortening of the manubrium with age. Days 20-30: N=11 medusae, 31-40: N=7, 41-50: N=25, 51-60: N=11, 61-70: N=34, 71-80: N=53. Some individual medusae were longitudinally measured at multiple ages. The *p*-value indicates the overall effect of age assessed using a linear mixed-effects model. (G) Schematic illustration of a *Cladonema* medusa tentacle. (H) Representative images of tentacles from young and aged medusae. Arrowheads indicate (ii) ocelli and (iii) nematocyte clusters. (I) Quantification of tentacle length showing age-associated shortening. Young: n=6 tentacles (N=3 medusae), Aged: n=9 (N=3). (J) Representative images of ãH2AX immunostaining in tentacle bulbs from young, middle-aged, and aged medusae, showing age-associated accumulation of DNA damage. Dashed boxes indicate the regions quantified in (K). Arrowheads indicate representative γH2AX^+^ cells, and insets show magnified views of representative cells. (K) Quantification of ãH2AX^+^ cells in tentacle bulbs. *p*-values shown are Holm-adjusted for multiple comparisons. Young: n=9 tentacles (N=3 medusae), Middle: n=9 (N=3), Aged: n=9 (N=3). Scale bars: (D, Hi) 1 mm, (Hii, Hiii) 200 µm, (J) 50 µm.

The hydrozoan jellyfish *Cladonema pacificum* has recently emerged as a tractable model for studying regeneration in medusae (Accorsi et al. 2024; Fujita et al. 2021). While retaining robust regenerative capacity during early adulthood, *Cladonema* medusae follow a reproducible aging trajectory marked by progressive tissue deterioration (Fig. 1B), providing a physiologically relevant window in which to examine how aging and regeneration intersect. Importantly, *Cladonema* tentacles, similar to other hydrozoan medusae such as *Clytia* (Leclère et al. 2012), harbor well-characterized stem and progenitor populations, including multipotent interstitial stem cells (i-cells) and regeneration-associated undifferentiated cells, which together allow branching morphogenesis, sustain tissue homeostasis, and drive rapid appendage regeneration (Hou et al. 2021; Fujiki et al. 2019; Fujita et al. 2019; Fujita et al. 2022; Fujita et al. 2023; Fujita et al. 2026). In line with efforts to broaden regeneration research beyond canonical systems, *Cladonema* offers practical advantages, including ease of laboratory maintenance, a short and observable life cycle, and experimentally accessible regenerative tissues. By focusing on the medusa stage, age-related changes in tissue homeostasis and regeneration can be examined within a single life-history context.

In this study, we systematically investigate how aging affects tissue homeostasis and regeneration in *Cladonema* medusae. Using tentacles as a tractable model tissue, we quantify age-dependent changes in differentiated cell maintenance, stem cell dynamics, and regenerative responses following injury. Our results show that aging disrupts regeneration in *Cladonema* through coordinated alterations in stem cell-associated processes, revealing aging as a conserved constraint on regeneration even in early-branching metazoans. Together, these findings establish *Cladonema* as an experimental model for studying aging–regeneration interactions and provide insight into how aging reshapes regenerative systems across animal evolution.

## Materials and Methods

### Animal cultures and surgical manipulations

*Cladonema pacificum* (female strain 6W) medusae were used for this research (Takeda et al. 2018). The medusae were maintained in plastic containers (V-type containers, V-7, V-8, V-9; AS ONE) at 22°C in artificial seawater (ASW) prepared using SEA LIFE (Marin Tech). Medusae of the same age were maintained together. The medusae were fed ad libitum with Vietnamese brine shrimp (Artemia; A&A Marine LLC) every 3-4 days. The seawater was replaced 1 h after feeding and again on the following day. Under standard culture conditions, medusae were subjected to a daily dark condition of 30 min for spawning, while being kept under constant light conditions during the remaining time (Masuda-Ozawa et al. 2022).

For quantification of egg production, the medusae were individually maintained in separate wells of a 6-well plate under the same culture conditions described above. Ten minutes after the end of the dark period on the day following feeding, the number of eggs released into each well was counted under a stereomicroscope. Egg production was repeatedly quantified in the same six medusae throughout the longitudinal observation period.

Before surgical manipulation, the medusae were anesthetized with 7% MgCl_2_ in deionized water for 2 min. Surgical manipulations were performed using micro scissors. For tentacle-based assays, three tentacles were amputated per medusa. After manipulations, the medusae were immediately returned to ASW. For tentacle amputation experiments, medusae were fed 1 day before manipulation and were starved after amputation, except for the 7 dpa prey-capture assays, in which medusae were additionally fed at 3 and 6 dpa.

Morphological observations were performed using a stereomicroscope. Bright field pictures of medusae were taken with a LEICA S8APO microscope with a Nikon digital camera (D5600). Longitudinal morphological observation was initiated at 20 days of age. Before imaging, medusae were transferred to a dish and relaxed by adding 7% MgCl_2_ at approximately a 1:5 ratio to the seawater.

For age-group comparisons, medusae ≤40 days old were classified as young, those between 41 and 60 days old as middle-aged, and those ≥61 days old as aged.

### Prey-capture assays

For the whole-medusa prey-capture assay (Fig. S2D and Fig. S2E), medusae were starved for 2 days before the assay. Individual medusae were placed in wells of a 24-well plate containing 2 mL of ASW and 20 Artemia. After 1 h, Artemia remaining free in the seawater and those attached to the tentacles were counted. The number of ingested Artemia was calculated by subtracting the numbers of free and tentacle-attached Artemia from the initial number of 20. Artemia were thus classified as ingested, captured (remaining attached to a tentacle), or no capture (remaining free in the seawater).

For the regenerated-tentacle prey-capture assay (Fig. 4D and 4E), medusae were examined at 7 dpa. Three regenerated tentacles were tested per medusa. An individual Artemia was held with forceps and brought into contact with each regenerated tentacle. The outcome was classified as ingested when the Artemia attached to the tentacle and was subsequently transported to the manubrium within the 1-min observation period (Thoma et al. 2023), captured when the Artemia remained attached but was not transported to the manubrium, or no capture when the Artemia failed to remain attached to the tentacle. For each medusa, the number of the three tested tentacles assigned to each category was recorded. Medusae that died before the assay were excluded from the analysis.

### Immunofluorescence

The *Cladonema* medusae were anesthetized with 7% MgCl_2_ in deionized water for 5 min and fixed for 1 h at room temperature (RT) or overnight at 4°C with 4% paraformaldehyde (PFA) in ASW. After fixation, the samples were washed 4 times (5 min each) with PBS containing 0.1% Triton X-100 (0.1% PBT). The samples were incubated with 3% H₂O₂ in PBS for 10 min, followed by 3 washes (5 min each) with 0.1% PBT. The samples were incubated overnight at 4°C with primary antibodies in 0.5% blocking buffer (0.5% blocking reagent [Roche] in Maleic acid buffer). The antibodies used were rabbit anti-phospho-Histone H2A.X (Ser139) (1:100; GeneTex, GTX127342), rabbit anti-FMRFamide (1:1000; ImmunoStar, 20091), mouse anti-β-catenin (8E4) (1:100; Enzo, ALX-804-260-C100), rabbit anti-phospho-Histone H3 (Ser10) (1:500; Upstate, 06–570), and rabbit anti-PKC ζ C-20 (aPKC; 1:100; Santa Cruz, sc-216). After primary antibody incubation, the samples were washed 4 times (10 min each) with 0.1% PBT and incubated in 0.5% blocking buffer for 1 h at RT. The samples were incubated overnight with Mouse MAX PO or Rabbit MAX PO (1:20; Nichirei Corporation), corresponding to the host species of the primary antibody, in 0.5% blocking buffer. The samples were washed 3 times (10 min each) with 0.1% PBT, followed by 3 washes (10 min each) in TNT buffer (100 mM NaCl, 100 mM Tris-HCl, pH 9.5, 0.1% Tween-20). The samples were incubated with either Cy3-tyramide solution (TSA Plus Cyanine 3; AKOYA Biosciences, NEL744001KT) or Cy5-tyramide solution (TSA Plus Cyanine 5; AKOYA Biosciences, NEL745001KT) for 10 min. The samples were washed 3 times (10 min each) with 0.1% PBT. Finally, the samples were incubated with Hoechst 33342 (1:500) in 0.1% PBT for 30 min in the dark and washed 3 times (10 min each) with 0.1% PBT. The samples were mounted on slides with 70% glycerol.

Nuclei and poly-γ-glutamate were stained with DAPI (1:250; Invitrogen, D1306), and actin fibers were stained with Alexa Fluor 488 or 546 phalloidin (1:400; Invitrogen, A12379 or A22283) in 0.1% PBT for 1 h. Confocal images were acquired using a Zeiss LSM 880 confocal microscope. Image processing and quantification were performed using ImageJ/Fiji software.

### EdU labeling

The *Cladonema* medusae were incubated with 20 µM 5-ethynyl-2’-deoxyuridine (EdU) (EdU kit; Invitrogen C10337) in ASW for 24 h. For analysis of proliferative activity during regeneration, EdU exposure was initiated immediately after tentacle amputation and continued for 24 h. After EdU treatment, the samples were anesthetized with 7% MgCl_2_ in deionized water for 5 min and fixed with 4% PFA in ASW for 60 min at RT or overnight at 4°C. After fixation, the samples were washed 3 times (5 min each) with 0.1% PBT and incubated with an EdU reaction cocktail (1×reaction buffer, CuSO_4_, Alexa Fluor 647 azide, and 1×reaction buffer additive; Invitrogen) for 30 min at RT in the dark. After the EdU reaction, the samples were washed 3 times (10 min each) with 0.1% PBT and incubated with Hoechst 33342 (1:500) in 0.1% PBT for 30 min in the dark with gentle agitation. The samples were then washed 3 times (10 min each) with 0.1% PBT and mounted on slides in 70% glycerol.

For combined EdU labeling and fluorescence *in situ* hybridization (FISH), FISH was performed after EdU incorporation and fixation, followed by the EdU detection reaction after tyramide signal development, as described previously (Fujita et al. 2023).

### Fluorescent *in situ* hybridization

Purified total RNA was reverse transcribed into cDNA by PrimeScript II 1st strand cDNA Synthesis Kit (Takara, 6210A). A cDNA library was used as a template for PCR (*Nanos1, Nanos2, Piwi* and *Mcol1*). The primer sets used for PCR cloning are as follows: *Nanos1*: 5′-AAGAGACACAGTCATTATCAAGCGA-3′ (forward) and 5′-AGCACGTAAAATTGGACACGTCG-3′ (reverse), *Nanos2*, 5’-ACTTCTCCAAAACCTCATGCCGAG-3’ (forward) and 5’-GAATGGCGGGCGATTTGACATCC-3’ (reverse); *Piwi*, 5’-CACACAAGAGTTGGACCGGA-3’ (forward) and 5’-ACCGGCTTATCGATGCAACA-3′ (reverse); *Mcol1*: 5′-CTCGTCGGTATTGCCCTCTC-3′ (forward) and 5′-CCAACCTATCGTGGACGTGT-3′ (reverse). PCR products were subcloned into the TAK101 vector (TOYOBO). The resulting plasmids were used for RNA probe synthesis with digoxigenin (DIG) labeling mix (Roche, 11277073910), and T7 (Roche, 10881767001) or T3 RNA polymerase (Roche, 11031163001) was used, according to the insert direction.

Medusae were anesthetized with 7% MgCl_2_ in deionized water for 5 min and fixed with 4% PFA overnight at 4°C. The samples were washed 3 times (10 min each) with PBS (prepared with DEPC-treated water) containing 0.1% Tween-20 (0.1% PBST). The samples were incubated in hybridization buffer (HB buffer; 5×SSC, 50% formamide, 0.1% Tween-20, 50 µg/mL tRNA, 50 µg/mL heparin) for 15 min, followed by pre-hybridization with replaced HB buffer for 2 h at 55°C. The samples were hybridized with HB buffer containing the antisense probes (final probe concentration: 0.5–1 ng/µL in HB buffer) overnight at 55°C. After hybridization, the samples were sequentially washed with buffer 1 (5×SSC, 50% formamide, 0.1% Tween-20), buffer 2 (2×SSC, 50% formamide, 0.1% Tween-20), and 2×SSC, each for 15 min twice at 55°C. The samples were then incubated with 0.1% PBST for 15 min at RT and incubated in 1% blocking buffer (1% blocking reagent [Roche] in Maleic acid buffer) for 1 h. The samples were then incubated with anti-DIG-POD antibodies (1:500; Roche, 11207733910) in 1% blocking buffer overnight at 4°C. The samples were washed 3 times (5 min each) in TNT buffer and incubated with Cy3-tyramide solution (TSA Plus Cyanine 3; AKOYA Biosciences, NEL744001KT) or Cy5-tyramide solution (TSA Plus Cyanine 5; AKOYA Biosciences, NEL745001KT) for 10 min. Finally, the samples were washed 3 times (10 min each) with 0.1% PBST and incubated with Hoechst 33342 (1:500) in 0.1% PBST for 30 min in the dark. The samples were washed 3 times (10 min each) with 0.1% PBST and mounted on slides with 70% glycerol.

For combined *Nanos1* FISH and aPKC immunostaining, FISH was first performed through Cy3-tyramide signal development. After three washes with 0.1% PBST, samples were incubated with 3% H_2_O_2_ for 10 min to inactivate residual HRP activity. Samples were then washed three times with 0.1% PBT and processed for immunostaining as described above, beginning with primary antibody incubation. The aPKC signal was developed using Cy5-tyramide solution.

### Paraffin embedding and sectioning

Medusae were anesthetized with 7% MgCl_2_ in deionized water for 5 min. For tentacle histology used for nematoblast scoring, the manubrium was removed, and the samples were trimmed to retain the umbrella margin with attached tentacles. The samples were then fixed with 4% PFA overnight at 4°C. After fixation, samples were washed 3 times (5 min each) with PBS, briefly rinsed in deionized water to remove salts, and dehydrated through an ethanol series (70%, 80%, and 95% ethanol for 30 min each, 100% ethanol for 30 min twice and 60 min once), and cleared in xylene 3 times (30 min each). The samples were infiltrated with paraffin at 70°C in four steps (pure paraffin overnight; fresh paraffin for 60 min twice; and fresh paraffin overnight), embedded in paraffin, and solidified at RT. Paraffin blocks were sectioned at a thickness of 5 µm using a sliding microtome, and sections were mounted on glass slides and stored at RT until staining.

### Hematoxylin and eosin staining of paraffin sections

Paraffin sections were deparaffinized in xylene three times (5 min each) and rehydrated through an ethanol series (100%, 90%, 80%, and 70% ethanol, 1 min each), followed by rinsing with tap water for 10 min.

The sections were stained with Mayer’s hematoxylin solution (Muto Pure Chemicals, 30002) for 10 min and washed with tap water for 10 min. The sections were then counterstained with eosin solution (1% eosin Y [Muto Pure Chemicals, 32002] diluted sixfold in 60% ethanol and adjusted to pH 4.8–5.0 with acetic acid) for 5 min, followed by washing with tap water for 10 min. The sections were dehydrated through an ethanol series (70%, 80%, 90%, and 100% ethanol), cleared in xylene three times (1 min each), and mounted with HSR mounting medium. Bright-field images were acquired using a BZ-X810 or BZ-X1000 microscope (Keyence).

For histological assessment of nematoblast-rich regions, one section was selected from each tentacle in which the tentacle bulb was most completely represented. The nematoblast-rich region of the bulb was scored on a four-point scale (0-3) based on both cell density and thickness of the cell-rich layer. Score 0 represented a sparse and thin cell layer; score 1, a moderately dense but thin layer; score 2, a moderately dense and relatively thick layer; and score 3, a densely populated and thick layer. Scoring was performed blinded to age group.

### Image analysis and quantification

Morphometric measurements were performed using the segmented line tool of ImageJ/Fiji. Umbrella size was measured in lateral-view images as the perpendicular distance from the apex of the umbrella to the umbrella margin. Manubrium size was measured from its junction with the umbrella to the oral end. Tentacle length was measured from the proximal-most end of the tentacle bulb to the tip of the main tentacle, excluding lateral branches. The elongation rate at each time point was calculated by normalizing the tentacle length to its length immediately after amputation (0 dpa). Three amputated tentacles were measured for each medusa, and the mean elongation rate of the three tentacles was used as the value for individual medusa. Age groups designated as 30, 40, 50, 60, or 70 day-old included medusae within ±3 day-old of the indicated age.

Manubrium saturation (Fig. S5B) was quantified as an index of manubrium condition. Bright-field color images were converted to hue–saturation–brightness (HSB) stacks using ImageJ/Fiji, and the mean saturation value was measured within a 200 µm x 200 µm square region of interest (ROI) positioned at the center of the manubrium. Images were acquired under identical imaging settings.

Cell numbers were quantified using ImageJ/Fiji. Signal-positive cells were manually counted using the multipoint tool. Total cell number quantification was performed as follows: (1) Binarize the signal of Hoechst staining using the “Threshold” command. (2) Separate adjacent nuclei using the “Watershed” command. (3) Measure the number of nuclei using the “Analyze Particles” command.

For quantification of FMRFamide-positive neural cell bodies at the tentacle base (Fig. 2F and 2G), the boundary between the umbrella and tentacle was traced using the segmented line tool of ImageJ/Fiji. A region extending 15 µm on either side of this line was defined as the quantification area, and FMRFamide-positive cell bodies within this region were counted manually.

**Figure 2.**
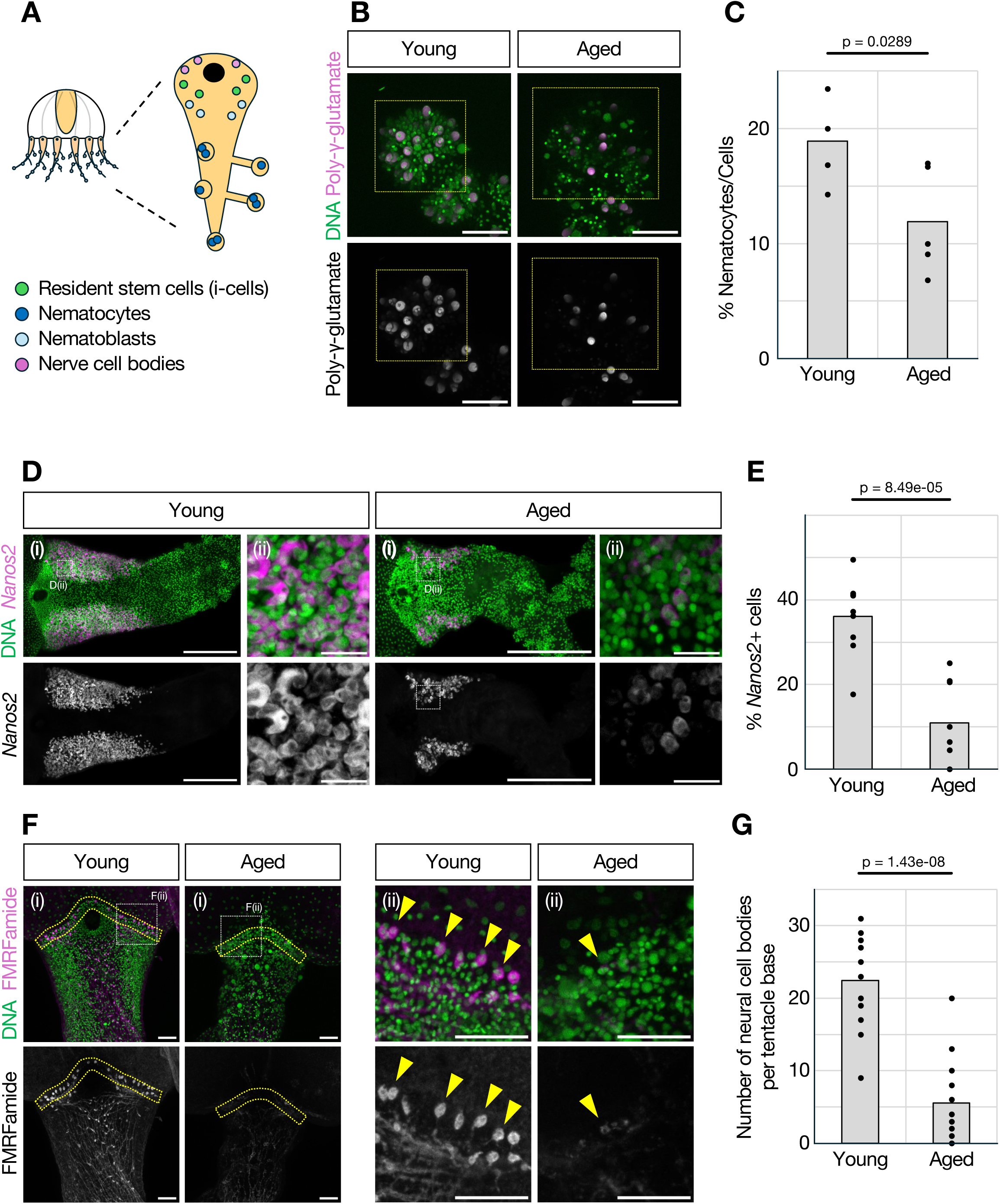
Differentiated cell populations in tentacles coordinately decrease with aging. (A) Schematic of the distribution of tentacle cell types. (B) Poly-ã-glutamate staining, a mature nematocyte marker, showing fewer mature nematocytes in aged tentacle tips. Dashed boxes indicate the region quantified in (C). (C) Quantification of mature nematocytes in tentacle-tip clusters. Young: n=5 tentacles (N=2 medusae), Aged: n=5 (N=3). (D) Fluorescent *in situ* hybridization (FISH) for *Nanos2*, a nematoblast marker, showing reduced numbers of *Nanos2*^+^ cells in aged tentacles. Dashed boxes indicate the region quantified in (E). Panels (ii) show magnified views of the boxed regions in (i). (E) Quantification of *Nanos2*^+^ cells in tentacle bulbs. Young: n=9 tentacles (N=3 medusae), Aged: n=8 (N=3). (F) FMRFamide staining, a neuronal marker, showing fewer neurons at tentacle bases in aged medusae. Yellow dashed regions indicate the regions quantified in (G). Panels (ii) show magnified views of the white boxed regions in (i). Arrowheads indicate neural cell bodies. (G) Quantification of neurons at tentacle bases. Young: n=15 tentacles (N=5 medusae), Aged: n=14 (N=5 medusae). Scale bars: (B, F) 50 µm, (Di) 200 µm, (Dii) 20 µm.

The extent of wound healing (Fig. S6A and S6B) was classified based on F-actin organization, following Fujita et al. (2023): “full open” when actin fibers at the wound remained unattached, “partially closed” when actin fibers were disorganized and closure was incomplete, and “completely closed” when actin fibers were fully attached across the wound (Fig. S6A).

Cell types in β-catenin– or aPKC-stained samples (Fig. 3B, 3C, 5E, 5F, S4A, and S4B) were classified based on staining patterns and nuclear morphology as described previously (Fujita et al. 2023).

**Figure 3.**
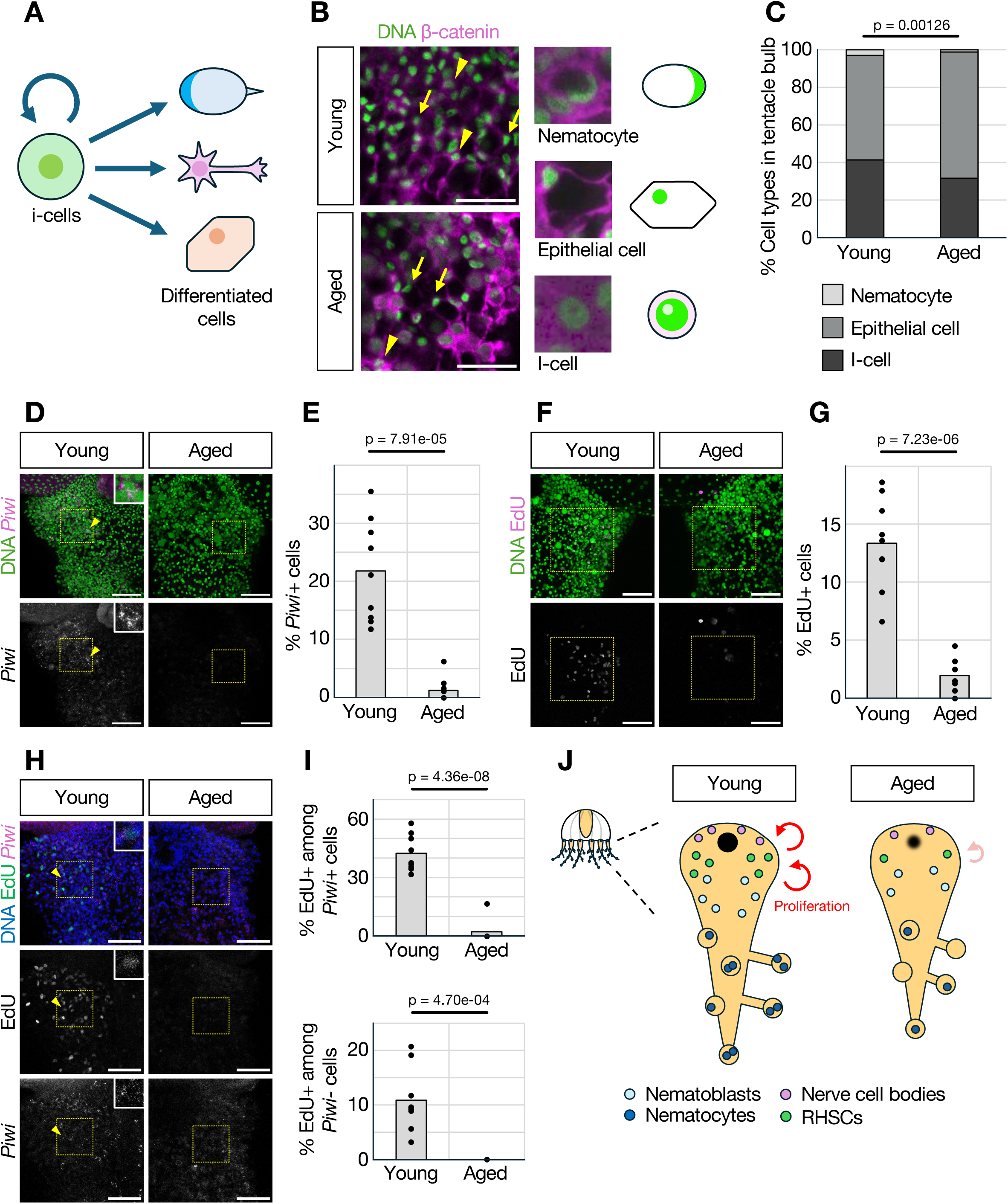
Resident stem cells in tentacles undergo functional decline with aging. (A) Scheme of stem cell (i-cell) maintenance and differentiation. (B) â-catenin staining showing altered cell-type proportions in tentacle bulbs, with fewer morphology-defined i-cell-like cells and a higher proportion of epithelial cells. Arrows indicate epithelial cells, and arrowheads indicate i-cell-like cells. (C) Quantification of cell-type proportions in tentacle bulbs based on â-catenin staining and cell morphology. Young: n=7 tentacles (N=3 medusae), Aged: n=5 (N=3). (D) FISH for *Piwi*, an i-cell marker, showing reduced numbers of *Piwi*^+^ cells in aged tentacle bulbs. Dashed boxes indicate the region quantified in (E). Arrowheads indicate representative *Piwi*^+^ cells, and insets show magnified views of representative cells. (E) Quantification of *Piwi*^+^ cells in tentacle bulbs. Young: n=9 tentacles (N=3 medusae), Aged: n=9 (N=3). (F) EdU staining showing fewer S-phase cells in aged tentacle bulbs. Dashed boxes indicate the region quantified in (G). (G) Quantification of EdU^+^ cells in tentacle bulbs. Young: n=9 tentacles (N=3 medusae), Aged: n=7 (N=3). (H) Combined EdU labeling and *Piwi* FISH in tentacle bulbs from young and aged medusae. Dashed boxes indicate the region quantified in (I). Arrowheads indicate representative EdU^+^/*Piwi*^+^ cells, and insets show magnified views of representative cells. (I) Quantification of the percentage of EdU^+^ cells among *Piwi*^+^ cells (upper) and among *Piwi*^-^ cells (lower). Young: n=9 tentacles (N=3 medusae), Aged: n=9 (N=3). (J) Schematic model showing the distribution of resident homeostatic stem cells (RHSCs), nematoblasts, nematocytes, and nerve cells in the tentacle. Scale bars: (B) 20 µm, (D, F, H) 50 µm.

**Figure 4.**
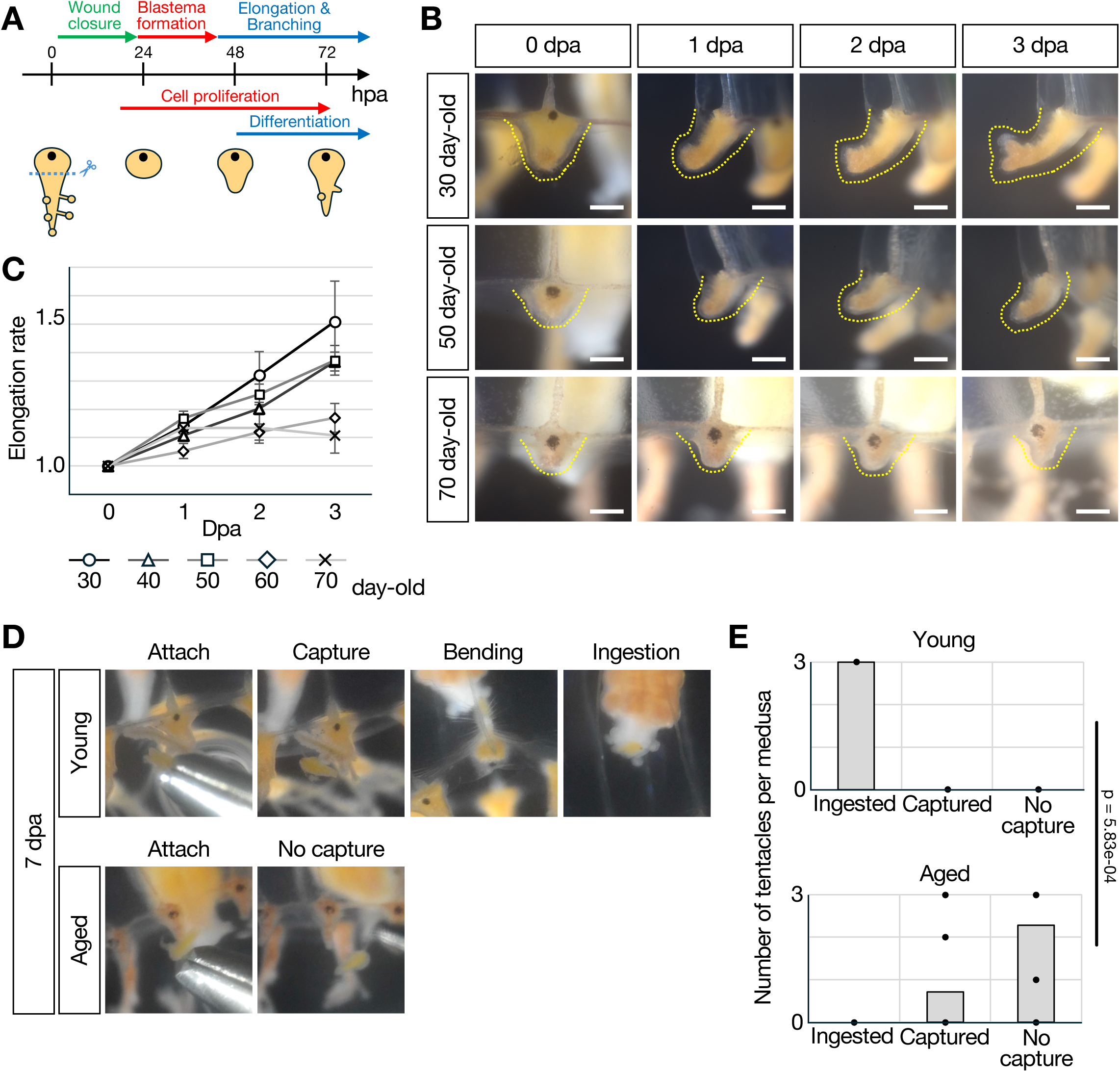
Regenerative elongation and functional recovery decline with aging. (A) Scheme of the tentacle regeneration process and its timeline. (B) Representative images of tentacle regeneration after amputation. (C) Elongation rate of regenerating tentacles from 0 to 3 dpa at different ages. 30 day-old: N=12 medusae, 40 day-old: N=16, 50 day-old: N=18, 60 day-old: N=10, 70 day-old: N=15. (D) Representative prey-capture and ingestion behaviors of regenerated tentacles at 7 dpa. (E) Quantification of prey-capture and ingestion outcomes of regenerated tentacles at 7 dpa. Each dot represents one medusa. Young: N=7 medusae, Aged: N=7. Scale bars: (B) 200 µm.

**Figure 5.**
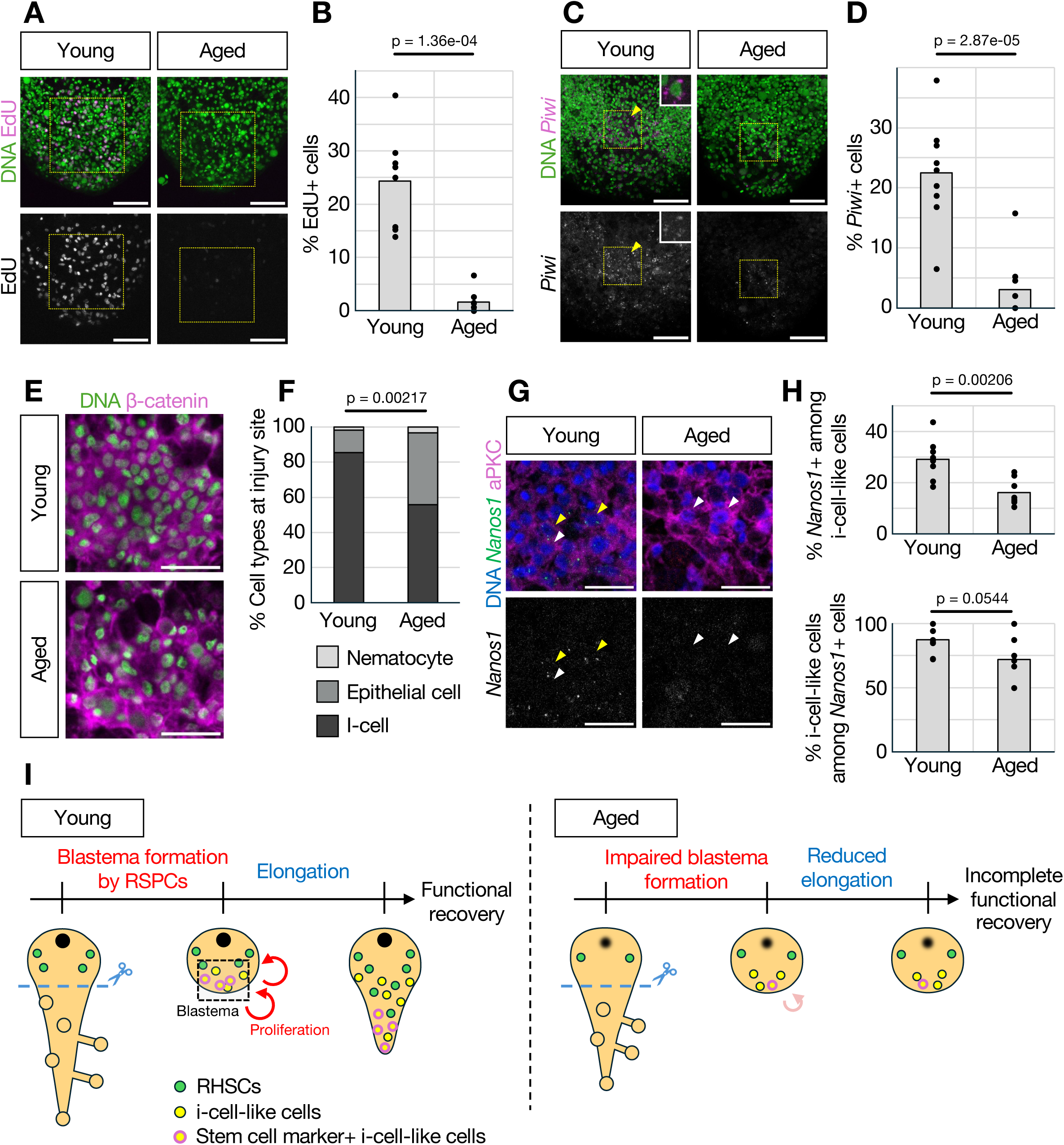
Blastema formation is impaired with aging. (A) EdU labeling showing reduced numbers of S-phase cells at injury sites in aged medusae. Dashed boxes indicate the regions quantified in (B). (B) Quantification of EdU^+^ cells at injury sites. Young: n=8 tentacles (N=3 medusae), Aged: n=8 (N=3). (C) FISH for *Piwi* showing reduced numbers of *Piwi*^+^ cells at aged injury sites. Dashed boxes indicate the regions quantified in (D). Arrowheads indicate representative *Piwi*^+^ cells, and insets show magnified views of representative cells. (D) Quantification of *Piwi^+^* cells at injury sites. Young: n=9 tentacles (N=3 medusae), Aged: n=9 (N=3). (E) â-catenin staining showing altered cell-type proportions at injury sites in aged medusae, with fewer morphology-defined i-cell-like cells and a higher proportion of epithelial cells. (F) Quantification of cell-type proportions at injury sites. Young: n=6 tentacles (N=3 medusae), Aged: n=6 (N=3). (G) Combined *Nanos1* FISH and aPKC immunostaining at injury sites from young and aged medusae. Yellow arrowheads indicate *Nanos1*^+^ i-cell-like cells, and white arrowheads indicate *Nanos1*^-^ i-cell-like cells. (H) Quantification of the relationship between *Nanos1* expression and i-cell-like morphology. The upper graph shows the percentage of *Nanos1*^+^ cells among morphology-defined i-cell-like cells. The lower graph shows the percentage of morphology-defined i-cell-like cells among *Nanos1*^+^ cells. Young: n=8 tentacles (N=3 medusae), Aged: n=8 (N=3). (I) Schematic model of age-associated impairment of blastema formation and tentacle regeneration. RHSCs, resident homeostatic stem cells; RSPCs, repair-specific proliferative cells. Scale bars: (A, C) 50 µm; (E, G) 20 µm.

For combined EdU and *Piwi* analyses (Fig. 3H and 3I), the proportion of EdU^+^ cells among *Piwi*^+^ cells was calculated as the number of EdU⁺/*Piwi*⁺ cells divided by the total number of *Piwi*⁺ cells. The proportion of EdU^+^ cells among *Piwi*^-^ cells was calculated as the number of EdU⁺/*Piwi*⁻ cells divided by the total number of *Piwi*⁻ cells.

For combined *Nanos1* FISH and aPKC staining (Fig. 5G and 5H), cells with i-cell-like morphology were classified based on aPKC staining and nuclear morphology. The proportion of *Nanos1*^+^ cells among morphology-defined i-cell-like cells and, conversely, the proportion of morphology-defined i-cell-like cells among *Nanos1*⁺ cells were calculated separately.

### Statistical analysis

For the tentacle elongation assay, the individual medusa was treated as the biological replicate and statistical unit, with measurements from three tentacles averaged to obtain one value per medusa.

Statistical analyses were performed using Microsoft Excel and R. Two-tailed Welch’s *t*-tests were used for comparisons between 2 groups. For analyses involving multiple pairwise comparisons within the same predefined family, *p*-values were adjusted using the Holm method. Age-dependent changes in umbrella size, relative manubrium size, and egg production (Fig. 1E, 1F, and S1C) were analyzed using linear mixed-effects models fitted by restricted maximum likelihood, with age as a categorical fixed effect and individual medusa as a random intercept; the overall effect of age was assessed using an F-test with Satterthwaite’s approximation. Categorical composition data from β-catenin– and aPKC-based cell-type classifications (Fig. 3C, 5F, and S4B) and prey-capture assays (Fig. 4E and S2E) were analyzed by PERMANOVA based on Bray–Curtis dissimilarities of within-sample category proportions; *p*-values were obtained by enumerating all unique group-label allocations. Nematoblast-rich area score distributions (Fig. S3E) were compared using the Fisher’s exact test. Associations between elongation rate at 3 dpa and morphological indices (Fig. S5C) were examined by multiple linear regression, with age group included as a categorical covariate; the *p*-value for each morphological index was reported as the age-adjusted *p*-value. Bar graphs show mean values, and dots represent individual values.

## Results

### Aging induces organism-wide morphological changes in *Cladonema* medusae

To characterize age-associated features in *Cladonema* medusae, we continuously monitored individuals under standardized culture conditions. *Cladonema* medusae possess three major organs: the bell (umbrella), which is essential for swimming; the manubrium, which mediates feeding and gametogenesis; and tentacles, which function in prey capture and defense against predators (Fig. 1C) (Fujiki et al. 2019; Fujita et al. 2019). We thus examined age-dependent changes in these organs over time (Fig. S1). As medusae aged, we observed pronounced reductions in umbrella size, regression of the manubrium, and shortening of tentacles (Fig. 1D). Among these changes, shrinkage of the umbrella and manubrium was particularly evident. Quantitative measurements confirmed that both organs undergo significant size reductions with age (Fig. 1E and 1F).

Based on these observation and measurements, we classified medusae with no obvious organ-level changes up to 40 days old as young. Medusae 61 days or older, which consistently exhibited reductions in umbrella and manubrium size, were classified as aged, whereas those between 41 and 60 days, which showed greater inter-individual variability, were classified as middle-aged. These reproducible, organism-wide morphological changes raised the question of how aging affects structural integrity and function at the tissue and cellular levels in *Cladonema* medusae.

### Manubrium and tentacle structural changes associated with aging

To examine age-dependent structural changes in greater detail, we performed histological analysis of paraffin sections stained with hematoxylin and eosin (H&E). In young medusae, the manubrium exhibited a well-organized bilayered structure composed of an outer ectodermal layer and an inner endodermal layer, with oocytes present in its internal region (Fig. S2A). In contrast, aged medusae showed marked thinning of the ectodermal layer, accompanied by an apparent reduction in the number of oocytes within the manubrium (Fig. S2A and S2B). These structural alterations suggest age-associated deterioration of reproductive tissues. To further assess age-related changes in reproductive output, we quantified egg production longitudinally after the onset of sexual maturity. The mean number of eggs released peaked during the young phase and subsequently declined with age (Fig. S2C), consistent with the structural changes observed in the aged manubrium (Fig. S2A).

We next examined tentacle morphology in detail. The tentacles of *Cladonema* medusae contain photoreceptor organs (ocelli) at their basal regions and clusters of nematocytes at their distal ends (Weber 1981; Fujita et al. 2023) (Fig. 1G). Ocelli in young medusae typically exhibited a regular, nearly circular morphology, whereas those in aged medusae frequently appeared irregular and disrupted (Fig. 1Hⅱ). Similarly, distal tentacles of young medusae harbored numerous well-defined spherical nematocyte clusters, whereas such clusters were less prominent in aged medusae (Fig. 1Hⅲ). Quantification further showed that tentacles were significantly shorter in aged than in young medusae (Fig. 1I). Together, these observations show that *Cladonema* medusae undergo reproducible, organism-wide morphological alterations with age.

To complement these morphological observations with a molecular indicator of age-associated cellular changes, we examined phosphorylated histone H2AX (γH2AX), a marker associated with DNA damage (Rogakou et al. 1998; Zhu et al. 2025). γH2AX-positive cells in the tentacle bulb increased with age and accumulated prominently in aged medusae (Fig. 1J and 1K). Thus, age-associated morphological decline was accompanied by the accumulation of a molecular marker of cellular damage. Based on these findings, we focused our subsequent analyses on the tentacles, in which cellular composition and regenerative processes have been relatively well characterized, to investigate cellular mechanisms underlying age-associated morphological and functional changes.

### Nematocyte-lineage and neuronal cell populations in tentacles decline with aging

Hydrozoan i-cells are multipotent stem cells that generate progenitors and differentiated cell types, including nematocytes and neurons. In both *Clytia* and *Cladonema* medusae, resident i-cells in the tentacles are concentrated near the tentacle base (Denker et al. 2008; Chari et al. 2021; Fujita et al. 2023) (Fig. 2A). To investigate cellular changes associated with tentacle regression, we compared nematocyte-lineage and neuronal populations in young and aged medusae.

Because distal nematocyte clusters became less prominent with age (Fig. 1Hiii), we first examined mature nematocytes, which are essential for prey capture, by staining poly-γ-glutamate synthesized in these cells (Szczepanek et al. 2002). Quantification revealed a significant reduction in the number of mature nematocytes in aged medusae (Fig. 2B and 2C). To assess the functional changes associated with aging, we performed a whole-medusa prey-capture assay (Fig. S2D). Nearly all *Artemia* presented to young medusae were ingested. In aged medusae, ingestion was substantially reduced, although many *Artemia* remained captured by the tentacles, indicating that prey-capture ability was at least partially retained (Fig. S2E). Thus, overall feeding performance declined with age, although this decline cannot be attributed solely to changes in the tentacles; regression of the manubrium and changes in prey transport or feeding behavior may also contribute.

Mature nematocytes in hydrozoan medusa tentacles arise from nematoblast precursors derived from i-cells (Denker et al. 2008; Fujita et al. 2023). We thus assessed nematoblast populations by fluorescence *in situ* hybridization (FISH) for *Mcol1*, an established nematoblast marker (Kanska & Frank 2013; Bradshaw et al. 2015; Flici et al. 2017; Fujita et al. 2023), and *Nanos2*, which is co-expressed with *Mcol1* in *Cladonema* nematoblasts (Fujita et al. 2023). In young medusae, *Nanos2*– and *Mcol1*-positive cells were broadly distributed from the tentacle bulb to the mid-region of the tentacle, whereas the numbers of cells positive for either marker were significantly reduced in aged medusae (Fig. 2D, 2E, S3A and S3B). Because these changes could reflect either reduced nematoblast abundance or age-dependent changes in marker expression, we also performed a marker-independent histological assessment.

Consistent with the FISH results, H&E-stained tentacle bulbs showed an age-associated shift toward lower scores for the density and thickness of the nematoblast-rich region (Fig. S3C–S3E). Together, these independent analyses support an age-associated reduction of the nematoblast lineage.

Neurons in cnidarians are likewise generated from i-cell lineages, diverging from shared stem cell populations into distinct neural progenitors (Chari et al. 2021). Age-related neuronal loss has been reported in other cnidarians, such as *Hydra oligactis* following the induction of sexual reproduction (Tomczyk et al. 2019). Using anti-FMRFamide immunostaining, we quantified neuronal cell bodies concentrated near the tentacle–umbrella junction. Compared with young medusae, aged medusae exhibited a significant reduction in neuronal number (Fig. 2F and 2G). These findings indicate that tentacle regression during aging is accompanied by reductions in both nematocyte-lineage and neuronal cell populations.

### Tentacle stem-like cells exhibit age-dependent alterations

In *Cladonema* medusae tentacles, as in those of other hydrozoan medusae, stem cell populations, or i-cells, are localized near the base and function as resident homeostatic stem cells (RHSCs), supplying progenitors and differentiated cell types required for tissue homeostasis (Denker et al. 2008; Fujita et al. 2023) (Fig. 3A). Given the age-associated loss of progenitors and differentiated cell populations, we next asked whether RHSCs themselves undergo age-associated changes.

To assess RHSCs abundance, we performed β-catenin immunostaining combined with nuclear morphology analysis (Fig. 3B). Hydrozoan i-cells were identified by cytoplasmic β-catenin localization, a high nuclear-to-cytoplasmic ratio, and prominent nucleoli (Plickert et al. 2012; Fujita et al. 2023). In contrast, epithelial cells displayed polygonal shapes with membrane-localized β-catenin, whereas nematocytes were distinguished by crescent-shaped nuclei distorted by nematocyst formation. Quantification revealed a modest reduction in the proportion of morphology-defined i-cell-like cells and a corresponding increase in epithelial cells in aged medusae (Fig. 3C). A similar shift was observed using atypical protein kinase C (aPKC) immunostaining and the same morphology-based classification criteria (Fujita et al. 2023) (Fig. S4A and S4B). Consistent with these observations, FISH for the stem cell-associated marker *Piwi* showed a marked reduction in *Piwi*-positive cells in aged medusae (Fig. 3D and 3E).

To evaluate proliferative activity in the tentacle bulb, we performed EdU incorporation assays, which revealed a pronounced decline in DNA synthesis in aged medusae (Fig. 3F and 3G). Similarly, immunostaining for the mitotic marker phospho-histone H3 (PH3) showed a significant reduction in mitotic cells with age (Fig. S4C and S4D). To identify the populations affected by this proliferative decline, we combined EdU labeling with *Piwi* FISH and analyzed EdU incorporation separately in *Piwi*-positive and *Piwi*-negative cells. The proportions of EdU-positive cells were significantly lower in aged medusae in both populations (Fig. 3H and 3I). Thus, the age-associated decline in proliferation was not restricted to *Piwi*-positive stem-like cells but extended to the broader *Piwi*-negative population. Moreover, total cell density in the tentacle bulb showed a marked reduction (Fig. S4E and S4F), suggesting defects in cell turnover.

Together, these results show that aging is associated with reductions in both marker-positive and morphology-defined stem-like cell populations, accompanied by a broader decline in proliferative activity within the tentacle bulb (Fig. 3J).

### Aging reduces regenerative capacity of *Cladonema* tentacles

We next asked whether aging affects regenerative capacity in *Cladonema* medusae. Even in highly regenerative organisms, such as axolotls, regenerative ability has been reported to decline with age (Faisal et al. 2024). Following amputation, young *Cladonema* tentacles normally undergo wound closure and blastema formation during the first day, begin nematogenesis and functional recovery by approximately 2 days post-amputation (dpa), and develop branches, nematocyte clusters, and prey-capture ability by approximately 3 dpa, although elongation toward the pre-amputation length may continue thereafter (Fujita et al. 2023) (Fig. 4A).

To determine how this regenerative response changes with age, we amputated tentacles from 30-, 40-, 50-, 60-, and 70-day-old medusae and monitored regrowth over time. Tentacles in young medusae elongated steadily and formed branches, whereas regeneration became less robust in older medusae (Fig. 4B, 4C, and S5A). This decline was not uniform across age groups but was particularly pronounced between 50 and 60 day-old (Fig. 4C and S5A).

We next examined whether individual condition could account for the decline in regeneration. After adjustment for age, umbrella size, relative manubrium size, and manubrium saturation were not significantly associated with tentacle elongation at 3 dpa (Fig. S5B and S5C). Thus, these morphological indices alone did not explain the age-associated decline in regeneration.

To assess longer-term functional recovery, we extended the analysis to 7 dpa and performed a prey-capture assay. All regenerated tentacles examined in young medusae captured and ingested prey, whereas most aged tentacles failed to capture prey, and none completed ingestion (Fig. 4D and 4E), suggesting that functional regeneration remained incomplete in aged medusae at 7 dpa.

During early regeneration, wound closure typically involves extensive remodeling of the actin cytoskeleton (Livshits et al. 2017). In young medusae, F-actin accumulated and reorganized at the wound margin as wound epithelial closure progressed, with most amputated tentacles achieving complete closure within 2 dpa (Fig. S6A and S6B). In aged medusae, however, this process was substantially delayed, and few tentacles achieved complete closure even by 3 dpa (Fig. S6A and S6B). Together, these results show that aging impairs tentacle regeneration from early wound closure through longer-term functional recovery.

### Aging impairs blastema formation

Functional tentacle regeneration in *Cladonema* requires formation of a blastema, a localized mass of proliferative, undifferentiated cells at the wound site (Aztekin 2021). Previous work showed that blastema formation in *Cladonema* involves repair-specific proliferating cells (RSPCs), which are largely distinct from RHSCs and preferentially contribute to epithelial cells during regeneration (Fujita et al. 2023). Systemic inhibition of cell proliferation by X-ray irradiation or pharmacological treatment, as well as local cell-cycle inhibition of RSPCs by UV laser exposure, reduces tentacle regeneration, indicating that proliferative activity, including that of RSPCs, is required for normal regeneration (Fujita et al. 2023). Given the reduced regenerative capacity of aged medusae, we hypothesized that blastema formation is compromised with age.

To test this, we examined proliferative activity at wound sites during regeneration. EdU incorporation assays revealed robust proliferation in the blastema of young medusae, whereas proliferative activity at wound sites was severely reduced in aged medusae (Fig. 5A and 5B). Similarly, PH3 immunostaining showed significantly fewer mitotic cells during regeneration in aged individuals (Fig. S6C and S6D).

Blastema cells in young medusae exhibit stem cell–like properties, including expression of the stem cell markers *Piwi* and *Nanos1* (Fujita et al. 2023). FISH revealed significantly fewer *Piwi*– and *Nanos1*-positive cells at wound sites in aged medusae (Fig. 5C, 5D, S6E and S6F). Similarly, β-catenin immunostaining combined with nuclear morphology showed a significant reduction in morphology-defined i-cell-like cells near aged wound sites (Fig. 5E and 5F). Combined *Nanos1* FISH and aPKC immunostaining further showed that most *Nanos1*-positive cells exhibited i-cell-like morphology but represented only a subset of broader morphology-defined i-cell-like populations (Fig. 5G and 5H). Moreover, the proportion of morphology-defined i-cell-like cells expressing *Nanos1* was significantly lower in aged medusae (Fig. 5H). Thus, marker-positive and morphology-defined stem-like populations overlap but are not equivalent. Collectively, these findings show that aging impairs blastema formation, as evidenced by reduced proliferation and diminished accumulation of blastema-associated stem-like cells.

## Discussion

In this study, we show that aging profoundly disrupts tissue homeostasis and regeneration in the cnidarian medusa *Cladonema pacificum*. By systematically analyzing age-dependent changes in morphology, cellular composition, and regenerative responses, we find that aging is accompanied by progressive morphological deterioration and accumulation of DNA damage, together with coordinated declines in differentiated cell maintenance, proliferative activity, and blastema formation (Fig. 3J and 5I). These defects are not confined to a single cellular compartment; rather, they impact multiple components of the regenerative system, including stem cell-associated populations in the tentacle bulb and injury-induced proliferative responses. Our findings together identify aging as a key constraint on blastema-mediated regeneration in *Cladonema* and show that even in early-branching metazoans with robust regenerative capacity, aging compromises the cellular programs that sustain tissue renewal and repair. Although regenerative capacity was not significantly associated with the morphological indices examined after adjustment for age (Fig. S5B and S5C), other age-associated systemic changes, including nutritional or physiological status, may also contribute to regenerative decline. Thus, the specific systemic factors linking organismal aging to regenerative impairment remain to be determined.

### Stem cell dynamics and potential niche changes associated with aging

Our results suggest that aging-associated regenerative decline in *Cladonema* medusae involves not only alterations in stem cell-associated dynamics but also potential changes in the local cellular environment that supports these cells (Fig. 3J and 5I). In tentacles of aged medusae, the reduction in proliferative cells within the tentacle bulb (Fig. 3F and S4C) and the decreased maintenance of progenitors and differentiated cell populations (Fig. 2B, 2D and 2F) indicate compromised stem cell–supported tissue turnover. In addition to reduced baseline proliferation, the attenuated injury-induced proliferative response observed in aged animals suggests impaired activation of regenerative programs (Fig. 5A, 5B, S6C, and S6D). Such defects may reflect intrinsic changes in stem or progenitor cells but could also arise from age-dependent alterations in the stem cell niche that normally provides instructive cues for proliferation and regeneration (Brunet et al. 2023). Given that tentacle regeneration in *Cladonema* relies on coordinated interactions between stem cells, progenitor states, and surrounding tissues (Fujita et al. 2023), aging-related changes in tissue architecture or signaling environments may further limit the transition from homeostatic maintenance to regeneration-specific cellular programs. Altogether, these observations support a model in which aging constrains regeneration through combined effects on stem cell dynamics and their supporting niche, rather than through the failure of a single cellular component.

### Blastema failure as an integrated outcome of aging

An important question raised by our findings is whether defective blastema formation represents a primary cause of regenerative failure in aged *Cladonema* medusae or instead reflects downstream consequences of aging-associated defects. Our data support the latter interpretation, in which blastema failure emerges as an integrated outcome of compromised stem cell dynamics and altered tissue environments (Fig. 3D, 3E, 5C, and 5D). In aged animals, reductions in baseline proliferation, attenuated injury-induced proliferative responses, and impaired maintenance of differentiated cells precede or accompany defects in blastema formation (Fig. 2B, 2C, 2F, 2G, 3F, 3G, 5A and 5B). This suggests that the blastema does not fail in isolation but rather reflects an inability of aging tissues to mobilize and coordinate key regenerative processes. Notably, this impairment cannot be explained solely by depletion of stem-like cells. Although a substantial population of morphology-defined stem-like cells remains at injury sites in aged medusae (Fig. 5E and 5F), cells expressing the stem cell markers *Piwi* and *Nanos1*, as well as proliferative activity, are markedly reduced (Fig. 5A-5D, Fig. S6C-S6F). This discrepancy suggests that aging may impair the ability of the remaining stem-like cells to enter or maintain the proliferative and molecular states associated with blastema formation (Fig. 5G and 5H). Whether this represents an exhaustion-like state or another form of age-associated functional decline will require direct lineage-tracing and long-term functional analyses. Consistent with this view, blastema formation can be considered a sensitive readout of regenerative system integrity (Seifert & Muneoka 2018).

Of note, previous work has shown that during tentacle regeneration in *Cladonema* medusae, RSPCs arise from a lineage distinct from RHSCs (Fujita et al. 2023). This lineage distinction suggests that blastema formation relies on regeneration-specific cellular programs: aging may therefore impair regeneration by selectively compromising the activation, expansion, or coordination of these regeneration-specific cell populations, potentially through age-associated changes in cellular state or the local tissue environment.

### Comparison with aging-associated regenerative decline in bilaterians

The aging-associated regenerative defects observed in *Cladonema* medusae share notable conceptual parallels with findings from bilaterian models that exhibit relatively high regeneration ability, including vertebrates. For example, in the short-lived African turquoise killifish, aging has been shown to impair regeneration by disrupting progenitor cell activation, proliferative capacity, and early regenerative responses in tissues including brain and fins (Wendler et al. 2015; Van houcke et al. 2021). Similarly, our analyses indicate that aging in *Cladonema* compromises both homeostatic cell turnover and injury-induced proliferation (Fig. 3F, 3G, 5A and 5B), leading to defective blastema formation. Despite profound differences in tissue organization and regenerative strategies between cnidarians and bilaterians, these observations suggest that aging commonly targets core components of regenerative systems, including stem or progenitor cell dynamics and their coordination with surrounding tissues.

Importantly, while bilaterian regeneration often involves inflammation and complex immune responses mediated by mobile immune cells and cytokine signaling (Mescher 2017; Godwin et al. 2013; Petrie et al. 2014), cnidarians appear to lack dedicated mobile immune cells and do not mount classical inflammatory responses. Nevertheless, cnidarians possess conserved innate immune signaling components, including pattern recognition receptors and downstream signaling pathways that mediate epithelial responses to microbial challenge and tissue perturbation (Emery et al. 2021; Wenger et al. 2014).

Whether age-dependent changes in such innate immune signaling contribute to regenerative decline in medusa-stage cnidarians remains an open question requiring further study. Together, these comparisons support the idea that aging imposes conserved constraints on regeneration across metazoans, while the cellular manifestations of this constraint remain context dependent.

### Aging–regeneration relationships across highly regenerative animals

Our findings in *Cladonema* provide a useful comparative framework for understanding how aging interacts with regeneration across animals with high intrinsic regenerative capacity. In cnidarians such as *Hydra*, continuous stem cell self-renewal and stable tissue organization support lifelong regeneration and negligible senescence under prolonged asexual conditions (Martínez 1998; Schaible et al. 2015). However, this apparent resistance to aging is not universal: in *Hydra oligactis*, induction of sexual reproduction is followed by functional deterioration, altered cell composition, and increased mortality, indicating that aging can emerge depending on life-history state and reproductive mode (Yoshida et al. 2006; Sun et al. 2020).

Similarly, planarian flatworms provide a complementary perspective in which strong regenerative capacity does not necessarily preclude aging. While some planarian lineages exhibit little evidence of aging under specific conditions (Mouton et al. 2011), recent work in the sexual lineage of *Schmidtea mediterranea* has demonstrated clear age-associated decline, together with the striking ability of regeneration to reverse multiple aging-related phenotypes (Dai et al. 2025). These findings indicate that regeneration can, in certain contexts, function as a rejuvenation mechanism rather than merely a repair process.

Collectively, these comparative examples highlight that strong regenerative capacity does not uniformly protect against aging-associated decline. Instead, aging–regeneration relationships appear to be shaped by the balance between stem cell maintenance, niche support, life-history state, and systemic regulation. In this context, *Cladonema* medusa represents a distinct scenario in which aging constrains both tissue homeostasis and regenerative responses within a finite lifespan, underscoring that regenerative potential alone is insufficient to counteract aging.

## Conclusion

In conclusion, our study establishes *Cladonema* medusae as a new experimental system for investigating how aging intersects with tissue homeostasis and regeneration. We show that aging disrupts both stem cell-supported tissue maintenance and blastema-mediated regeneration, indicating that robust regenerative capacity does not inherently confer resistance to aging-associated decline. These findings support the idea that aging represents a broadly shared constraint on regenerative systems and highlight how age-dependent alterations in stem cell dynamics and regenerative responsiveness shape regeneration outcomes across metazoans.

## Author contributions

R.K.: performed most experiments, analyzed data and prepared figures and manuscripts. H.N.: performed and supervised experiments. S.T.: performed and supervised histological analyses and edited manuscript. T.T.: supervised experiments and edited manuscripts. M.M.: supervised project and edited manuscript. Y.N.: supervised the whole project and experiments and wrote and edited manuscript.

## Acknowledgements

We are grateful to Aika Akahori and Dr. Taeko Kimura (Laboratory of Neuropathology and Neuroscience, Graduate School of Pharmaceutical Sciences,

The University of Tokyo) for technical guidance on paraffin embedding and sectioning.

## Data Availability Statement

Data available on request from the authors.

## Figure legends

**Figure S1.** Age-associated morphological changes across multiple individual medusae. Representative images of multiple *Cladonema* medusae at different ages, illustrating reproducible age-associated changes in overall body size and organ morphology across individuals. Scale bars: 1 mm.

**Figure S2.** Age-associated structural and functional changes in medusae. (A) Representative longitudinal paraffin sections of the manubrium from young and aged medusae stained with hematoxylin and eosin. (ii) and (iii) show magnified views of the aboral and oral regions indicated in (i). The arrowhead indicates an oocyte. (B) Representative transverse paraffin sections of the manubrium from young and aged medusae stained with hematoxylin and eosin. Panels (ii) show magnified views of the boxed regions in (i). (C) Average number of spawned eggs at the indicated ages. N=6 medusae. The *p*-value indicates the overall effect of age assessed using a linear mixed-effects model. (D) Schematic of the prey-capture assay. Twenty Artemia were provided to each medusa for 1 h. (E) Proportions of Artemia classified as Ingested, Captured, or No capture after 1 h. Young: N=11 medusae, Aged: N=12. Scale bars: (Ai, Bi) 200 µm; (Aii, Aiii, Bii) 50 µm.

**Figure S3.** Independent analyses confirm an age-associated reduction in nematoblasts. (A) Fluorescent *in situ* hybridization (FISH) for *Mcol1,* a nematoblast marker, showing reduced numbers of *Mcol1^+^* cells in aged tentacles. Dashed boxes indicate the regions quantified in (B). (B) Quantification of *Mcol1*^+^ cells in tentacle bulbs. Young: n=9 tentacles (N=3 medusae), Aged: n=9 (N=3). (C) Hematoxylin and eosin staining of paraffin sections of the tentacle from young and aged medusae. Arrowheads indicate nematoblast-like cells. Panels (ii) show magnified views of representative nematoblast-like cells indicated in (i). (D) Representative hematoxylin and eosin-stained sections illustrating the criteria for nematoblast-rich area scores from 0 to 3. Red dashed lines delineate the nematoblast-rich areas evaluated for scoring. (E) Distribution of nematoblast-rich area scores in young and aged tentacles. Young: n=16 tentacles (N=6 medusae), Aged: n=15 (N=5). Scale bars: (A) 50 µm; (Ci) 200 µm; (Cii) 10 µm; (D) 100 µm.

**Figure S4.** Age-associated cellular changes in tentacle bulbs. (A) aPKC staining showing altered cell-type proportions in aged tentacle bulbs, with fewer morphology-defined i-cell-like cells and a higher proportion of epithelial cells. (B) Quantification of cell-type proportions in the tentacle bulbs based on aPKC staining and cell morphology. Young: n=5 tentacles (N=3 medusae), Aged: n=5 (N=3). (C) PH3 staining showing fewer M-phase cells in aged tentacle bulbs. Dashed boxes indicate the regions quantified in (D). (D) Quantification of PH3^+^ cells in tentacle bulbs. Young: n=6 tentacles (N=3 medusae), Aged: n=5 (N=3). (E) Nuclear staining used to quantify total cell numbers in tentacle bulbs from young and aged medusae. Dashed boxes indicate the regions quantified in (F). (F) Quantification of total cell numbers within the defined regions of tentacle bulbs. Young: n=9 tentacles (N=3 medusae), Aged: n=8 (N=3). Scale bars: (A) 20 µm; (C, E) 50 µm.

**Figure S5.** Associations between regenerative elongation and age-associated morphological indices. (A) Elongation rate at 3 dpa in medusae of different ages. *p*-values shown are Holm-adjusted for multiple comparisons. 30 day-old: N=12 medusae, 40 day-old: N=16, 50 day-old: N=18, 60 day-old: N=10, 70 day-old: N=15. (B) Representative measurements of morphological indices used for the analyses, including umbrella size, manubrium size, and manubrium saturation. The red squares indicate the region used to quantify manubrium saturation. (C) Relationships between elongation rate at 3 dpa and umbrella size (left), the ratio of manubrium size to umbrella size (middle), and manubrium saturation (right) in 30-, 50-, and 70-day-old medusae. Age-adjusted *p*-values are indicated in each graph. 30 day-old: N=12 medusae, 50 day-old: N=12, 70 day-old: N=12.

**Figure S6.** Early wound healing and blastema-associated cellular responses are impaired with aging. (A) F-actin staining showing the process of wound healing from 0 to 3 dpa. F-actin was visualized with phalloidin. F, P, and C denote full open, partially closed, and completely closed, respectively. Yellow dashed outlines indicate the wound margins. (B) Quantification of the extent of wound healing during tentacle regeneration. Young, 0 dpa: n=9 tentacles (N=5 medusae), 1 dpa: n=9 (N=5), 2 dpa: n=9 (N=5), 3 dpa: n=9 (N=5); Aged, 0 dpa: n=9 (N=5), 1 dpa: n=7 (N=4), 2 dpa: n=7 (N=4), 3 dpa: n=7 (N=4). (C) PH3 staining showing fewer M-phase cells at injury sites in aged medusae. Dashed boxes indicate the region quantified in (D). (D) Quantification of PH3^+^ cells at injury sites. Young: n=8 tentacles (N=3 medusae), Aged: n=5 (N=3). (E) *Nanos1* FISH showing reduced numbers of *Nanos1*^+^ cells at aged injury sites. Dashed boxes indicate the regions quantified in (F). Arrowheads indicate representative *Nanos1*^+^ cells, and insets show magnified views of representative cells. (F) Quantification of *Nanos1*^+^ cells at injury sites. Young: n=9 tentacles (N=3), Aged: n=9 (N=3). Scale bars: (A) 100 µm; (C, E) 50 µm.

