## Supplementary Figures for "Aging disrupts tissue homeostasis and constrains blastema-mediated regeneration in the *Cladonema* medusa"

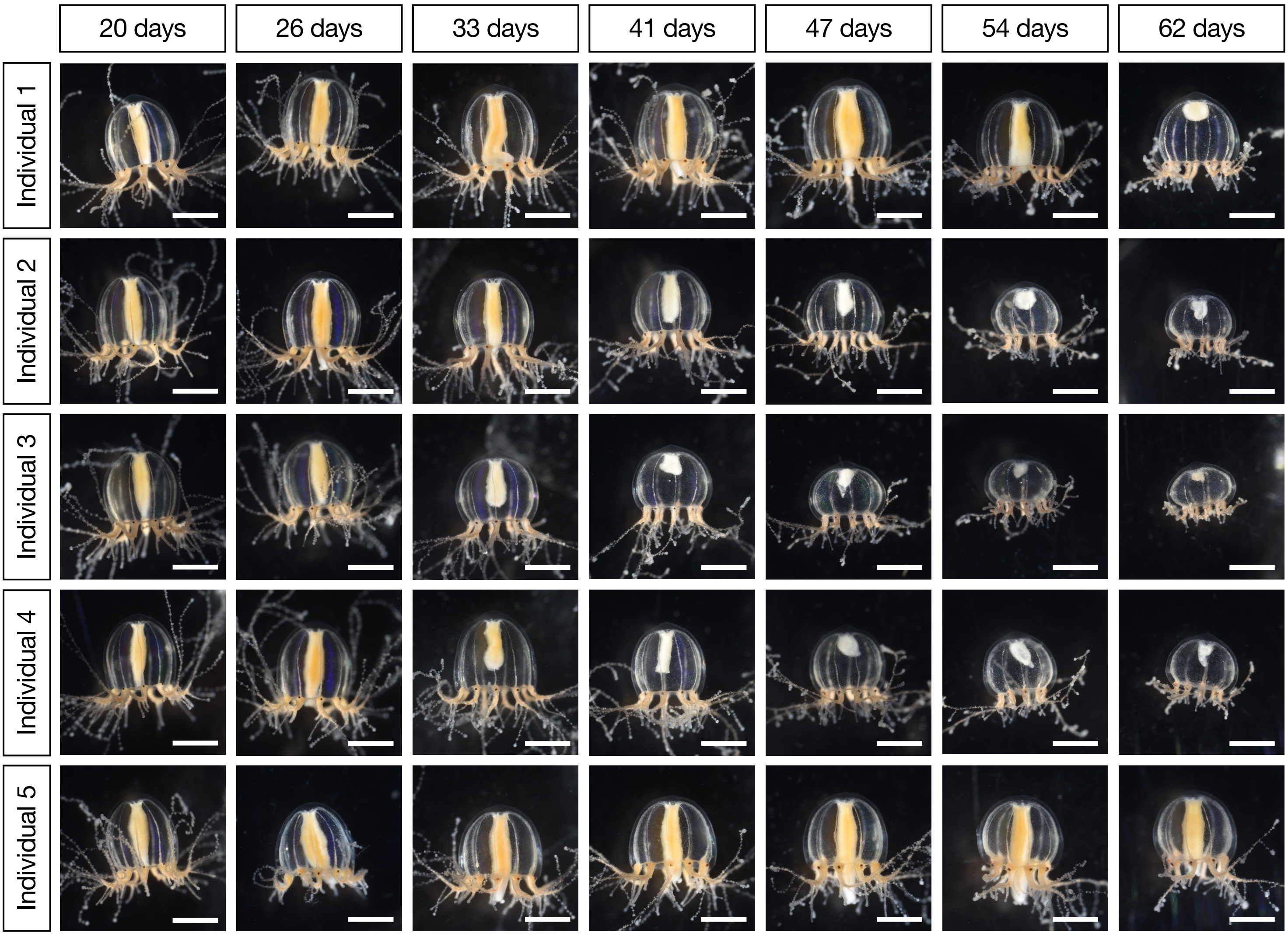


**Kanehisa et al.**

**Figure S1**

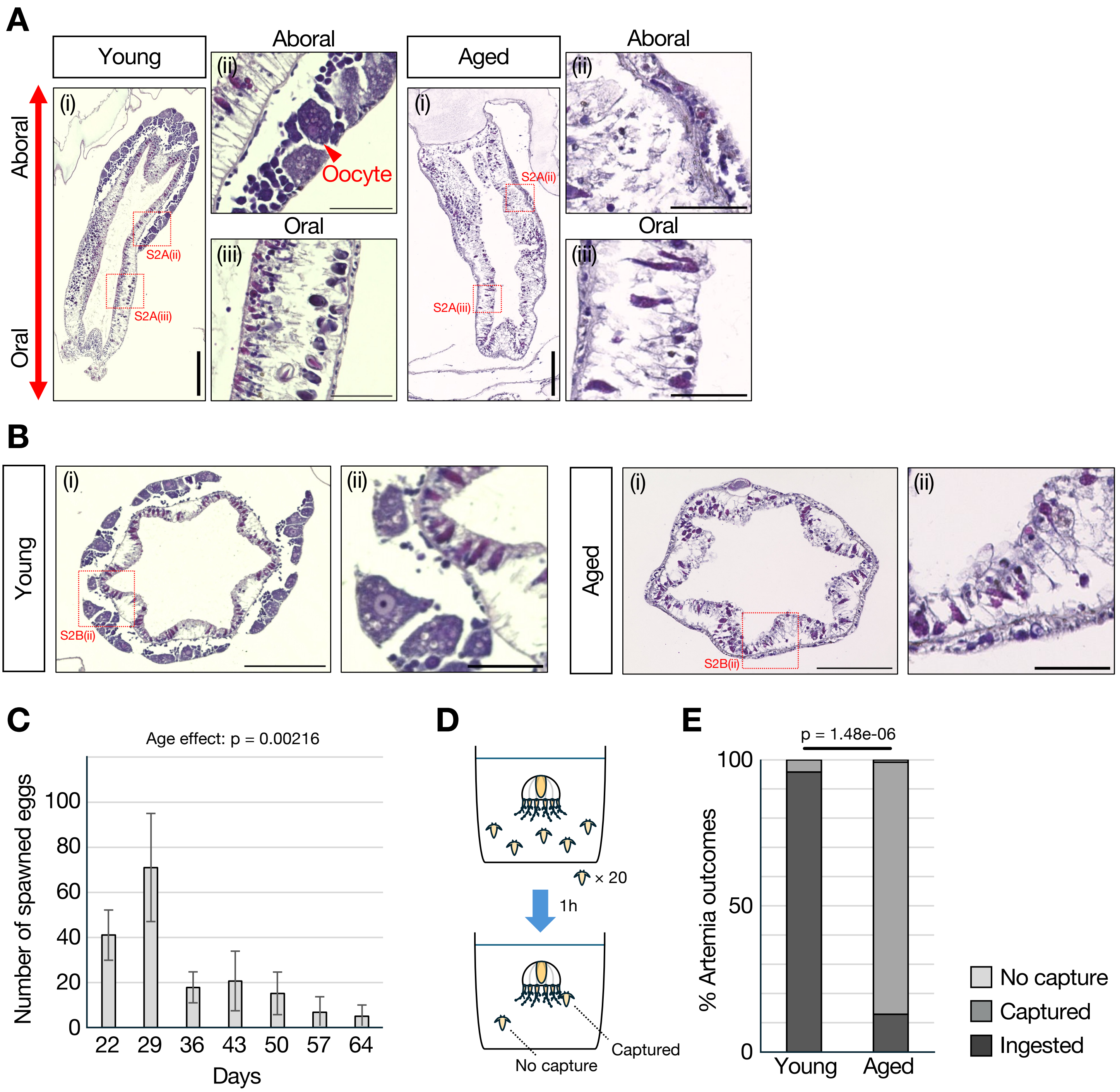


Kanehisa et al.

Figure S2

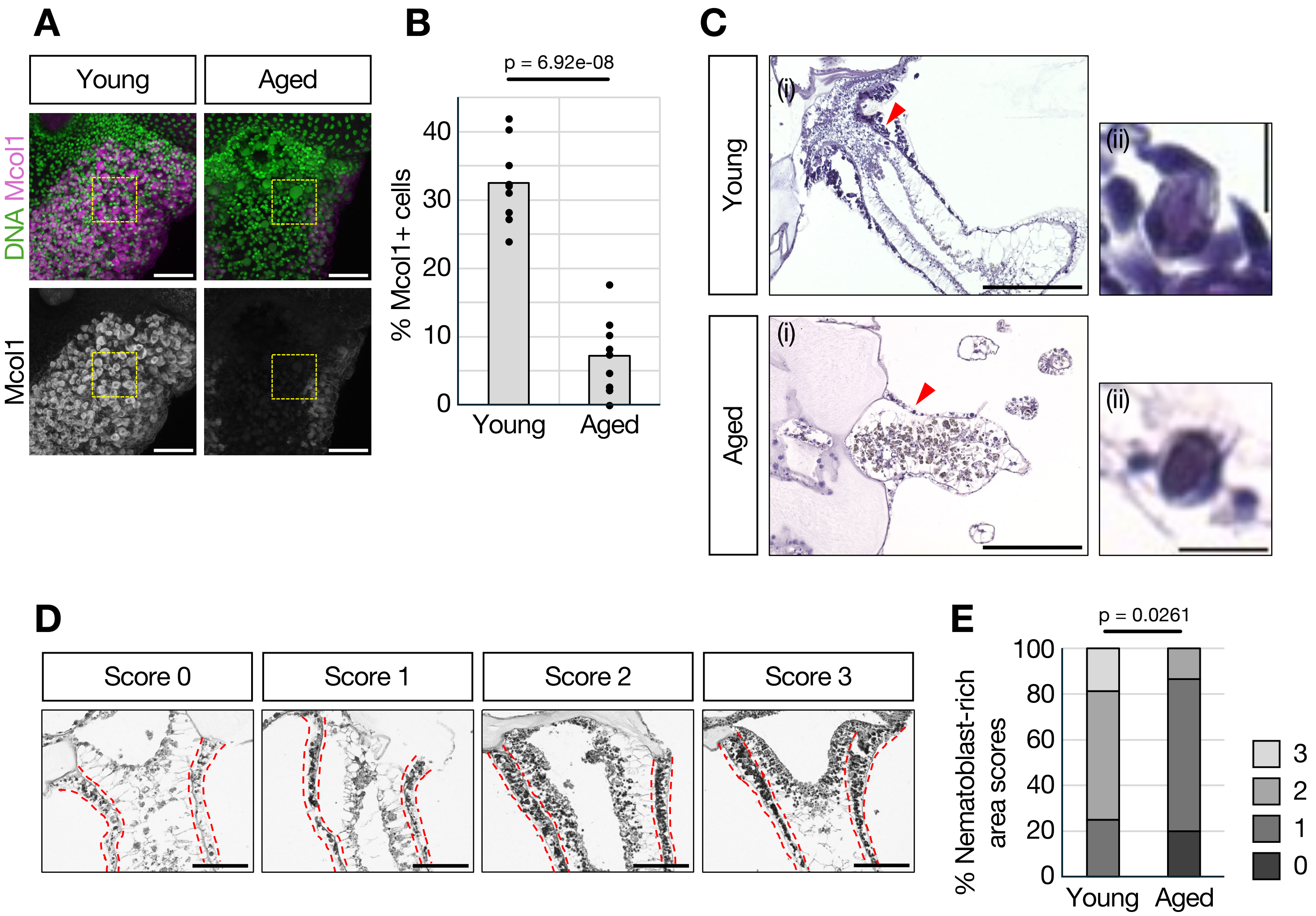


Kanehisa et al. Figure S3

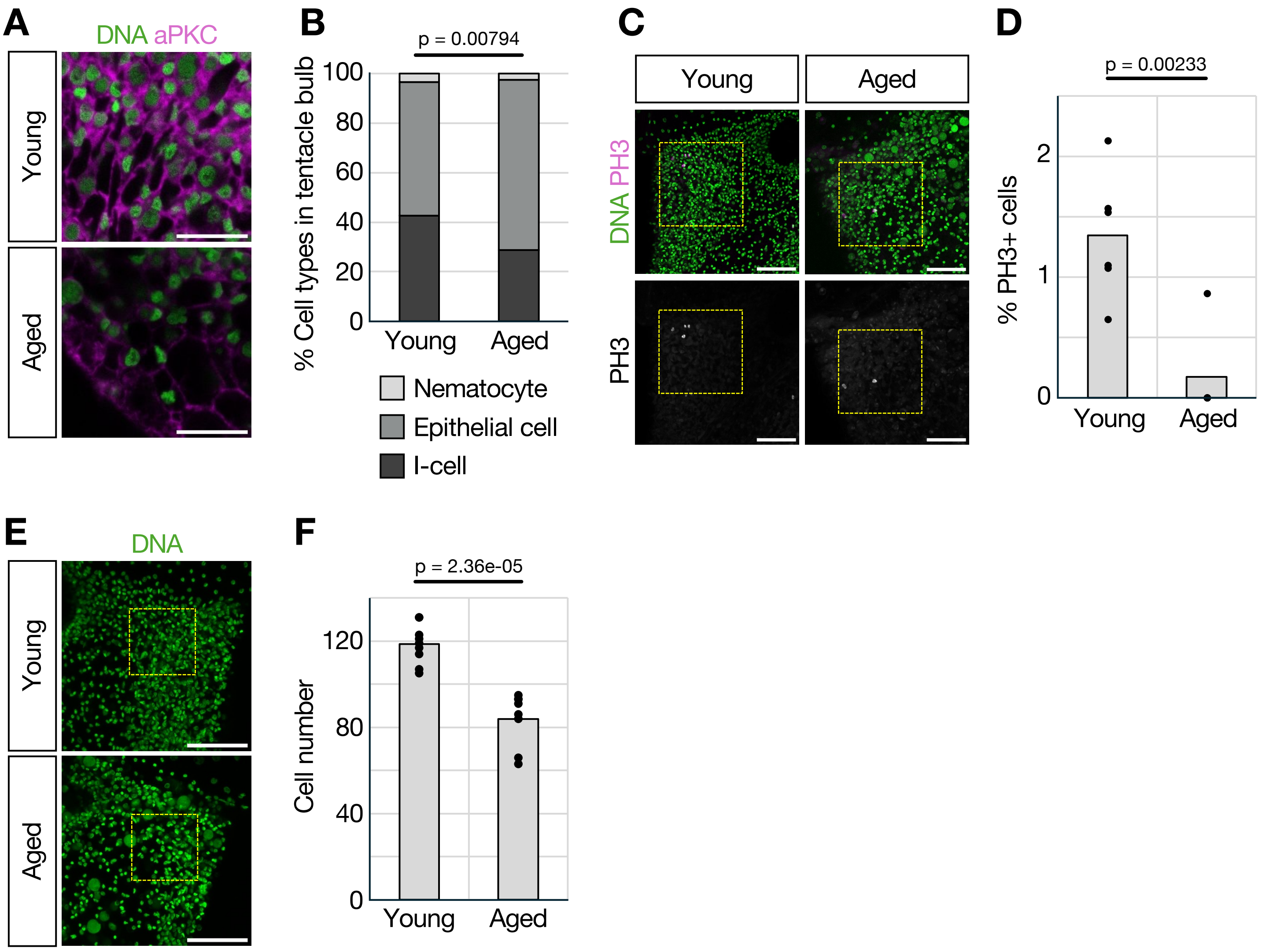


Kanehisa et al.

Figure S4

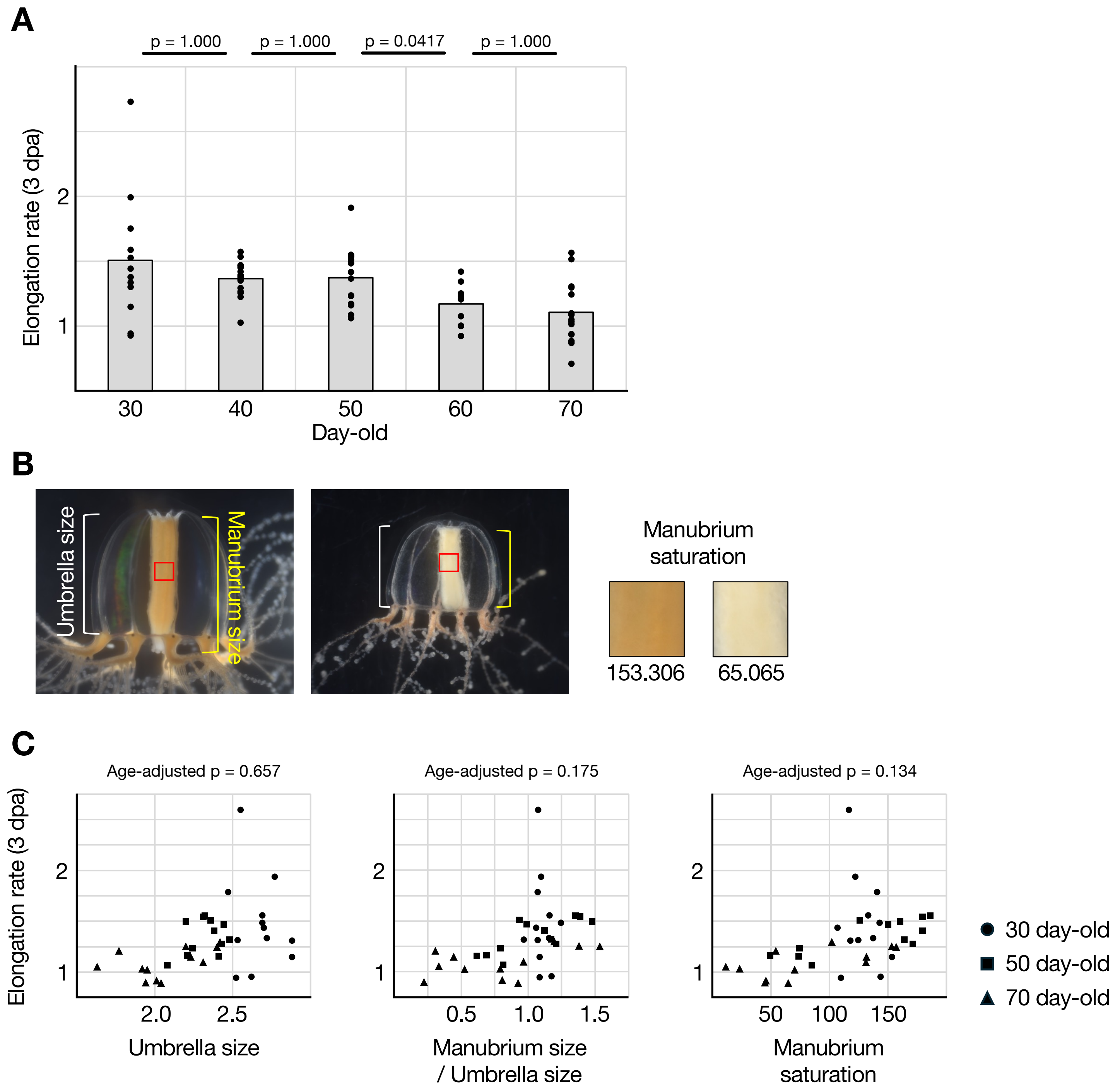


**Kanehisa et al.** **Figure S5**

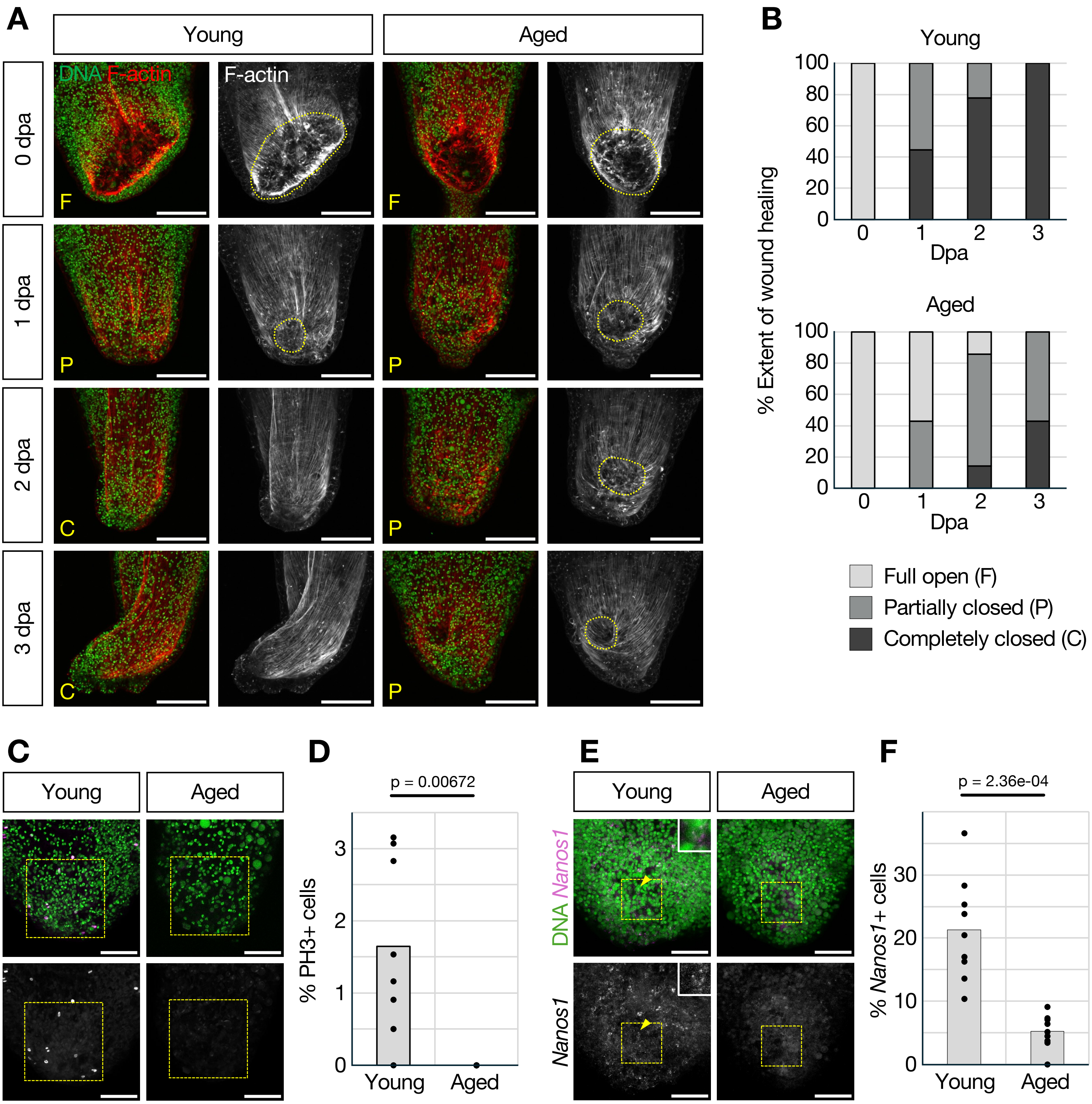


Kanehisa et al.

Figure S6
